# Observing Grazing Behavior Transitions in *Cafeteria roenbergensis* with Video-Rate Two-Photon Microscopy

**DOI:** 10.1101/2025.04.23.650267

**Authors:** Arifur Rahaman, Martin Chacon, Yuejiao Xian, Chuan Xiao, Chunqiang Li

## Abstract

Grazing behavior of free-living aquatic heterotrophic nanoflagellates (HNFs) on bacteria plays a central role in shaping microbial community structure and nutrient cycling. However, direct observation of these interactions has been limited by the rapid motility of flagellates and the transient nature of predator-prey encounters. To address this gap, this study presents a novel application of video-rate two-photon fluorescence microscopy for high resolution, real-time imaging of fast-moving microorganisms. HNF *Cafeteria roenbergensis* is used as a model system to investigate dynamic grazing interactions between fluorescently stained bacteria and the flagellates, where the flagellates are detected via their intrinsic cellular autofluorescence. This two-photon microscope combined with real-time imaging capability enables continuous capture of contact, capture, ingestion, and digestion steps during grazing. Quantitative analysis across varying prey concentration reveals phase-specific durations and saturation behavior in grazing activities. Furthermore, transitions in grazing dynamics across two continuous periods of flagellates in starved and fed conditions are observed in live time series videos. This technique provides a powerful new tool to study rapid microbial interactions *in situ* and can be broadly applicable to diverse microbe-microbe systems. With the integration of targeted fluorescent molecular probes, this technique holds significant potential for uncovering mechanical and biochemical interactions underlying microbial feeding and communication.

**Significance:** This study significantly advances microbial ecology by applying video-rate two-photon fluorescence microscopy to directly visualize and quantify the rapid grazing behavior of flagellates on bacteria. By enabling real-time observation of contact, ingestion, and digestion during the grazing events, the technique overcomes limitations of traditional imaging. It captures behavioral transitions tied to physiological states, offering quantitative insight into microbial predator-prey dynamics. This method establishes a broadly applicable platform for studying fast, transient microbial interactions. Although current limitations include the lack of 3D tracking, emerging optical technologies promise enhanced capabilities, paving the way for deeper understanding of microbial processes in aquatic ecosystems.

## Introduction

In most aquatic ecosystems, heterotrophic nanoflagellates (HNFs) are key bacterivorous organisms and play an important role in the microbial food web. Laboratory and field studies of interactions between HNFs and bacteria suggest that they are the main microbial consumers of bacteria and are able to regulate bacterial densities and diversity [1, 2]. One key research area is understanding the mechanisms of prey selection by HNFs. Advanced microscopy and imaging techniques can provide detailed visual information on these grazing processes thus elucidating mechanical and biochemical cues. Although the feeding ecology of several HNF species has been studied [3, 4], the predating behavior dynamics of HNFs has not been fully characterized. Fluorescence microscopy with different molecular probes can image subcellular organelles and provide molecular level information [5], and video-rate microscopy can record the grazing process in real time [3]. *Cafeteria roenbergensis*, a eukaryotic organism found in all oceans especially in coastal waters, is a common marine bacterivorous flagellate [6]. It has been used as a model system to study bacterial predation through observations of immobilized flagellates [3]. However, *C. roenbergensis* is a fast*-*moving HNF with a swimming speed of 10-100 µm/s. Current video microscopy techniques have only captured stationary predating patterns [3]. In addition, the expected interaction speed between *C. roenbergensis* and bacteria is also very fast, therefore, traditional imaging techniques face significant challenges in capturing their interactions.

Recently two-photon microscopy (TPM) has been widely used in biomedical research [7]. It is especially useful for exploring dynamic processes *in vivo* at cellular level [8]. The benefits of using two-photon excitation compared with conventional one-photon excitation include the intrinsic capability of optical sectioning at micrometer lateral resolution, deeper tissue penetration and reduced photobleaching and photodamage. By using fast scanning approaches, the imaging speed of TPM can achieve real-time video rate and even higher [9, 10]. Therefore, it is a suitable live imaging modality to monitor the interaction between fast moving HNF and bacteria. To observe the interaction between *C. roenbergensis* and bacteria, a home-built two-photon laser scanning fluorescence microscopy was developed. *C. roenbergensis* was imaged by two-photon excited autofluorescence in blue spectral region, and bacteria were stained using SYBR gold dye [11], then imaged simultaneously via two-photon excitation of SYBR Gold fluorescence at wavelength of 710 nm. By combing a polygonal mirror with a galvanometer to achieve 2D scanning, the imaging speed of the TPM system was configured to operate at video-rate (30 frames/s).

## Materials and Methods

### Preparation of C. roenbergensis and E. coli for Imaging

To prepare *C. roenbergensis* for imaging, 15 mL of culture is removed from a 125 mL flask and transferred to a centrifuge tube. The sample is centrifuged at 2,600× *g* for 20 minutes. After centrifugation, the supernatant is completely discarded, and the pellet is resuspended in 1 mL of 25 ppt of F/2 media. A second centrifugation is performed at 3,000× *g* for 20 minutes. Following this, the pellet is resuspended in 10-20 μL of 25 ppt F/2 media, depending on pellet size (10 μL for small, 20 μL for large). The cells are then stained with Lugol’s acid iodine solution and counted using a hemocytometer. Based on the cell count, the sample is diluted to the desired concentration. For *E. coli* preparation, 15 mL of culture is removed from main stock and centrifuge at 1,000× *g* for 1 minute to eliminate dead cells and aggregated cell chunks or debris. The top 8.5 mL of supernatant is transferred to a clean 15 mL tube. Optical density at 600 nm (OD_600_) is measured using a spectrophotometer and the bacteria suspension is diluted to a OD_600_ of 1.5, ensuring a final volume of at least 800 μL. A 1:200 dilution of SYBR Gold stain is prepared using 25 ppt F/2 media to obtain a final volume of 400 μL. Equal volumes (400 μL) of 1.5 OD_600_ *E. Coli* and the diluted SYBR Gold are mixed, resulting in a final concentration of 0.75 OD_600_ of *E. Coli* and 1:400 SYBR Gold. The mixture is incubated in the dark for 30 minutes. During incubation, an additional aliquot of 1.5 OD_600_ *E. Coli* is diluted with 25 ppt F/2 media to a final OD_600_ of 0.75 and a volume of 800 μL, serving as the unstained control. After incubation in dark, both stained and unstained samples are centrifuged at 2,000× *g* for 10 minutes. A volume of 700 μL of supernatant is removed from each sample and replace with 700 μL of 25 ppt F/2 media, followed by vortexing. This wash is repeated once more. The OD_600_ of the unstained sample is measured. Based on the OD_600_ reading, the stained sample again at 2,000 × *g* for 5 minutes. The pellet is then resuspended to reach a final OD_600_ of 1.2.

### Two-photon fluorescence imaging

Details of the custom-built two-photon microscope are described in the reference [10]. In summary, a mode-locked Ti/sapphire laser (Newport, Spectra-Physics Maitai HP, Wavelength 700-1020 nm, 100 fs, 80 MHz) is employed as the laser source; the laser beam is reflected to a fast-spinning polygonal mirror (480 revs/s) and a fast-swing galvanometer-mounted mirror (30 Hz) to achieve a 2D raster scan across the sample. A diachronic beam splitter (Semrock, FF660-FDi01) separates the excitation and emission beams; An Olympus 60× water immersion microscope objective lens (NA 1.2) focuses the laser beam onto the sample. The excitation wavelength is tunable via the Maitai HP control programmer on computer, and the excitation power on the sample can be adjusted using a combination of a quarter-wave plate and a linear polarizer as a laser attenuator. The blue detection channel, equipped with a band pass filters (Semrock FF02-447/60) to capture autofluorescence from *C. roenbergensis*. The green detection channel with a band pass filter (Semrock FF03-525/50) detects SYBR Gold fluorescence. A red detection channel with a band pass filters (Semrock FF01-593/46) is also built in the system. Fluorescence signal in each channel is detected by a photomultiplier tube (Hamamatsu R10699) and the electronic signals from three channels are acquired simultaneously using a frame grabber (Matrox Solios eA/XA). Videos are recorded on a personal computer at a rate of 30 frames/s. A reflection channel is implemented by inserting a beam splitter before the polygonal mirror. Reflected signals from the sample are collected by an avalanche photodiode (Hamamatsu S2381). A light source from a 660-680 nm laser diode hits on the polygonal mirror and is detected by a bi-cell to synchronize the polygonal mirror and galvanometer.

## Results and Discussion

### Imaging of HNF predator C. roenbergensis and prey bacteria

**Figure 1 (a)** shows the autofluorescence image of *C. roenbergensis* detected in the blue channel (420-470 nm) using excitation wavelength at 710 nm. The laser power on the sample was approximately 20 mW, and the photomultiplier tube (PMT) voltage was set to 3.0 V. For improved visualization, the original blue color was changed to the pseudo color red. **Figure 1 (b)** shows the green fluorescence image of bacteria stained with SYBR Gold, excited using the same laser beam as in **Figure 1 (a)** and detected in the green channel. The green fluorescence signal from the stained bacteria was significantly stronger than the autofluorescence signal from *C*. *roenbergensis*. While the bacteria were generally distributed uniformly across the frame, small clusters occasionally formed as indicated by the white circle (**Figure 1 (b)**). Live videos (30 frames/s) captured the rapid motility of *C. roenbergensis* and their dynamic interactions with bacteria in the surrounding (**Supplemental Video 1)**.

**Figure 1.**
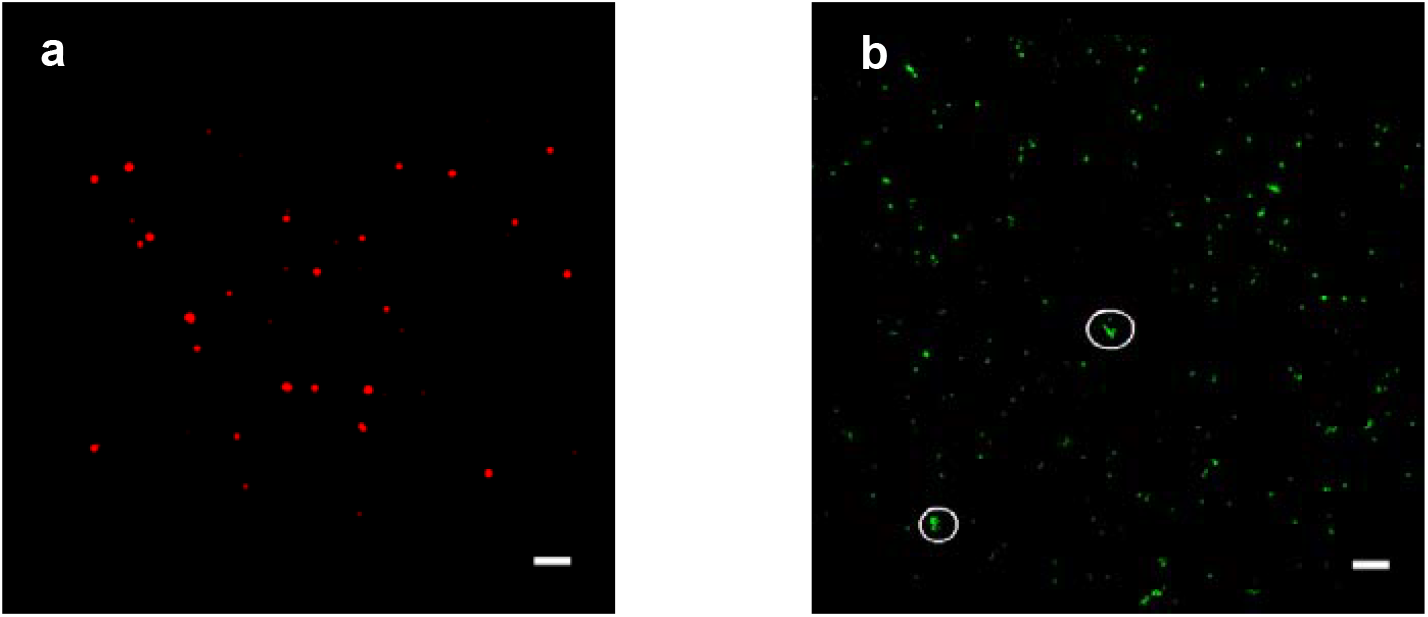
Fluorescent images of **(a)** *C. roenbergensis* in blue autofluorescence and **(b)** bacteria stained with SYBR Gold. Pseudo color red is used in **(a)** for improved visualization. (bar: 20 µm)

### Interaction between C. roenbergensis and bacteria

To visualize the interaction between *C. roenbergensis* and bacteria, 2 µL of each was mixed in a microcentrifuge tube. A 2-µL aliquot of this mixture was dripped onto a glass slide and covered with a cover glass. This prepared sample slide was immediately transferred under the two-photon microscope for imaging and recording. Typically, images and videos were acquired within 2 minutes after mixing up the samples. **Figure 2** presents a time series (2-60 minutes) of the interaction between *C. roenbergensis* and bacteria. Imaging was conducted using a laser power of approximately 20 mW on the sample. The PMT voltage was set to 1.5V and 3.0 V for the green and blue channel, respectively. In **Figure 2 (a)** to **(d)**, squares highlight the incidences just before *C. roenbergensis* engulfed bacteria, while circles indicate *C. roenbergensis* that have already engulfed the bacteria. When the two microbes overlap spatially, the merged fluorescence results in a yellow color. Over time, the number of overlapping particles increases, indicating active grazing behavior of *C. roenbergensis* on surrounding bacteria.

**Figure 2.**
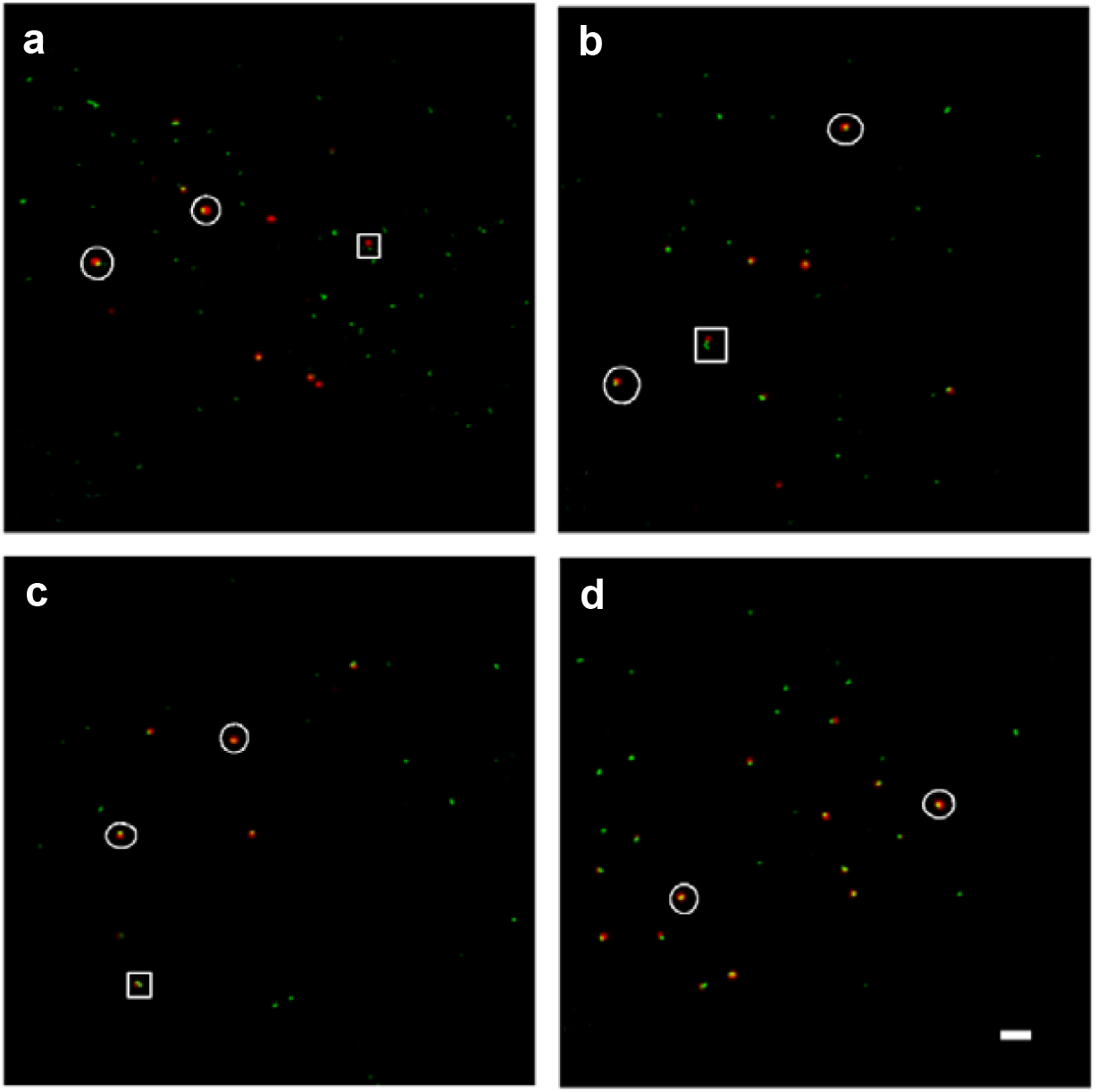
Interation between *C. roenbergensis* and bacteria **(a)** at 2 min after mixing up; **(b)** at 20 min after mixing up; **(c)** at 40 min after mixing up; **(d)** at 60 min after mixing up. Red: *C. roenbergensis*, Green: bacteria. (bar: 20 µm)

**Figure 3** shows sequential images from a time lapse video with 0.2 s time span. At the beginning, one fast swimming *C. roenbergensis* (red sphere in **Figure 3 (a)-(d)**) came into contact with one bacterium within 0.1 s. The engulfment of this bacterium happened within 0.03 s (**Figure 3 (e)** and **(f)**). At the end, this *C. roenbergensis* swam away with engulfed bacterium (**Figure 3 (g)**). Although the imaging speed of our two-photon microscope is fast enough to catch such events, due to the 3D sectioning capability of two-photon microscope, it is challenging to keep tracking the trajectory of *Cafeteria roenbergensis* when it moved out of focus in the z direction.

**Figure 3.**
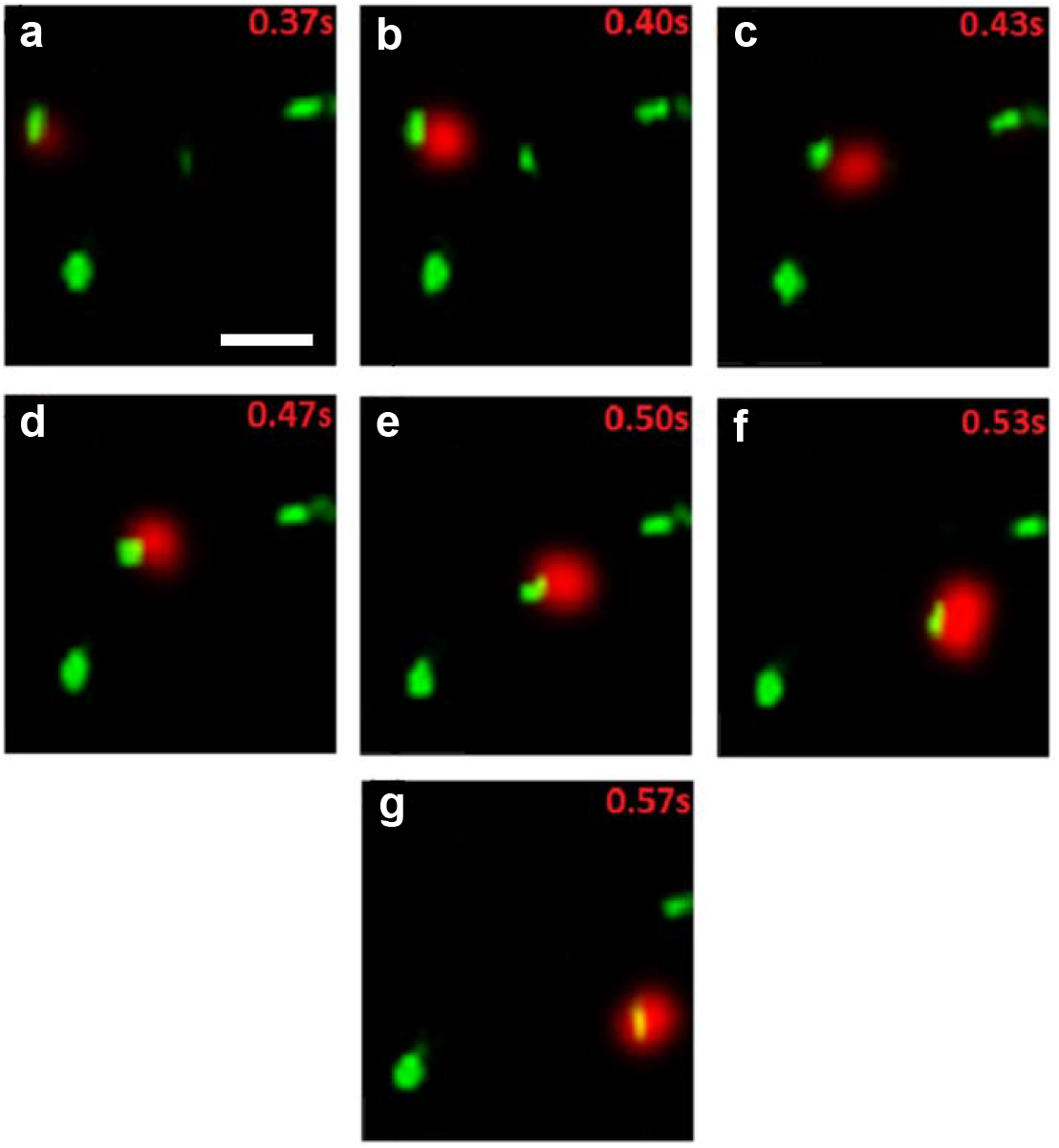
Predating behavior of *C. roenbergensis* on bacteria. Red: *C. roenbergensis*, Green: bacteria. (bar: 10 µm)

### Statistical analysis of interactions

Video-rate two-photon microscopy enables the direct observation of individual grazing interactions between *C. roenbergensis* and bacteria, making it advantageous for analyzing the dynamics of these events overtime. To reliably count the interaction events, noise reduction was performed separately on the blue and green fluorescence channels respectively due to the fact that the noise level in each channel is different. The green fluorescence from SYBR Gold stained bacteria is significantly stronger than the blue autofluorescence from *C. roenbergensis*. Particle counting was then performed on the denoised images to determine the number of *C. roenbergensis* particles, the bacteria particles, and overlapping particles (representing *C. roenbergensis* in contact with, engulfing, or ingesting bacteria). This experiment was repeated multiple times with three different concentrations of bacteria.

**Figure 4 (a)** plots the percentage of overlapping particles as time increases for three experiments. The total numbers of *C. roenbergensis* and bacteria in each experiment are listed in **Table 1**. It is obvious that in the first phase (0-9 minutes) the percentage of overlapping particles increases rapidly with a large slope. The initial slope (*V*_*0*_) is calculated using the first and second data points of this phase and listed in **Table 1**. After 9 minutes, this percentage increase reaches its saturation with a slope of about 0.5% per minute from simple estimation (second phase). After this saturation phase, the percentage increases again with a slope in the range of 1.3-2.3% per minute from three curves (third phase) and eventually it saturated at around 25 minutes and reached a steady state (fourth phase). These four phases can be divided into two periods (initial and later), each has a fast-increasing phase and a saturation phase. In each period, the two phases are consistent to an observation of starving flagellates have two phases of feeding. An initial phase for food vacuole formation and second phase for vacuole processing [4], where after 5-10 minutes the ingestion rate of starved *C. roenbergensis* significantly decreased [4]. In the first phase, *C. roenbergensis* ingests bacteria rapidly with an approximately linear rate. As bacteria being ingested by *C. roenbergensis*, digestion starts thus leading to the slower rate of grazing which shows up as the second phase. These kinetics can be mathematically described by the Michaelis-Menten equation commonly used to describe enzymes binding to substrates

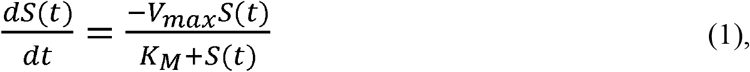

where *S(t)* is the substrate concentration, *V*_*max*_ is the maximum reaction rate, and *K*_*M*_ is the Michaelis constant [12, 13]. The amount of enzyme-substrate bound complex *ES(t)* formed is

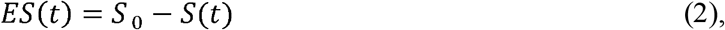

where *S*_*0*_ is the initial substrate concentration. The ordinary differential equation for *ES(t)* is obtained by combining equations (1) and (2) as

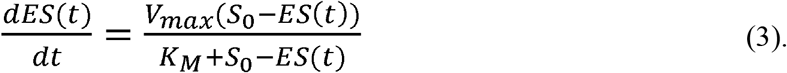

**Table 1.** Fitting parameters for the first and second phases (0-14 minutes)

| $C_0$ | $B_0$ | $V_0$ | $k_1 B_0$ | $k_1$ | $k_3$ | $K_M$ |
| --- | --- | --- | --- | --- | --- | --- |
| 46988 | 338997 | 25.2%/min | 0.158 | $4.66 \times 10^{-7}$ | 0.0306 | $6.44 \times 10^4$ |
| 26525 | 233488 | 16.1%/min | 0.127 | $5.44 \times 10^{-7}$ | 0.0691 | $1.27 \times 10^5$ |
| 20858 | 63641 | 12.8%/min | 0.106 | $1.67 \times 10^{-6}$ | 0.0892 | $5.33 \times 10^4$ |

**Figure 4.**
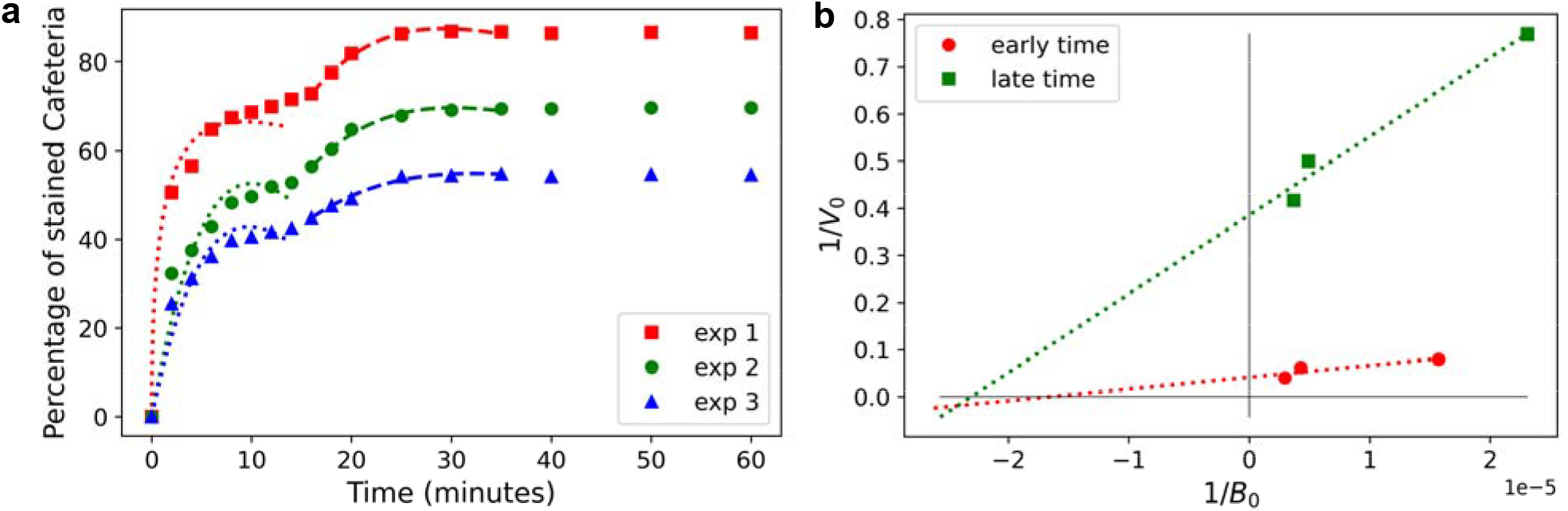
**(a)** Plot of percentage of overlapping particles vs. time. Three experiments are performed with different initial concentrations of *C. roenbergensis (C*_*0*_) and bacteria (*B*_*0*_) which are listed in **Table 1. (b)** Lineweaver-Burk plot of 1/*V*_*0*_ vs 1/*B*_*0*_

Hence, **Figure 4 (a)** is a progress curve of *ES(t) vs. t*. Integrating Eq. (3) will obtain an integral equation of *ES(t)* as

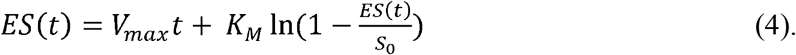

Analytically solving nonlinear differential equations (3) or (4) is impossible, therefore, regression methods have been developed to fit the experimental progress curve and obtain reaction parameters such as *V*_*max*_ and *K*_*M*_ [14, 15]. In this experiment *C. roenbergensis* act as enzymes and bacteria serve as the substrates, and the enzyme-substrate complex “ES” above is defined as those *C. roenbergensis* particles overlap with bacteria, denoted as “C*·*B*”* in the following. In order to have a more straightforward understanding of this progressive curve, instead of using the regression method we have developed a mathematical model (**Supplemental Methods**) to describe the time evolution of C*·*B particle numbers over total number of *C. roenbergensis* (C_0_) as

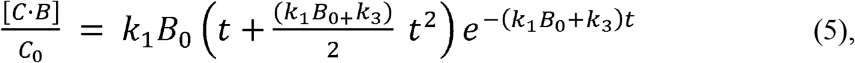

where [*C·B*] is the concentration of bacteria-overlapped *C. roenbergensis, C*_0_ is the total concentration of *C. roenbergensis, B*_0_ is the initial concentration of bacteria, *k*_*1*_ is the *C. roenbergensis* grazing rate, *k*_*3*_ is the digestion rate. Qualitatively the first term in bracket describes the fast growing of bacteria-overlapped *C. roenbergensis* concentration at initial time, and the exponential term describes the slowing down of this growth as time increases. Fitting experimental data with Eq. (5), parameters *k*_*1*_*B*_*0*_ and *k*_*3*_ will be obtained to estimate Michaelis *K*_*M*_ as *K*_*M*_ = *k*_3_/*k*_1_.

The dotted lines show the three fitting curves for each data set for the first and second phases only (0-14 minutes), and **Table 1** lists the fitting parameters for the three curves. Since it is difficult to estimate the *C. roenbergensis* and bacteria absolute concentrations due to the lack of precise measurement of the imaging volume, the total numbers of each type of particles observed during the experiment are used as *C*_0_ and *B*_0_. When the concentration of bacteria is higher, the grazing rate *k*_*1*_*B*_*0*_ is clearly higher. The fitted digestion rate *k*_*3*_ has large variations among three data sets. The calculated Michaelis constants *K*_M_ are shown in the last column. Although these estimates have large variations, they are in the same order, which demonstrates the qualitative validity of Eq. (5). In doing Taylor expansions to obtain Eq. (5) it is important to keep in mind that *k*_1_*B*_0_*t* and *k*_3_*t* need to be less than 1 for its validity. While obtaining these fitting results the period is within 14 minutes (*t*<14), thus showing *k*_1_*B*_0_*t* is slightly larger than 1 when the time is 10 minutes or later. This may explain the imperfection of these fittings, especially the fitting curves bend down when the *t* is large which is caused by the exponential term in Eq. (5).

The dashed lines show the three fitting curves for each data set for the third and fourth phases only (16-35 minutes, later period), and **Table 2** lists the fitting parameters for the three curves. In this table the number of bacteria *B*_*0*_ is modified since a significant number of bacteria have been ingested by *C. roenbergensis*. The modified bacteria *B*_*0*_ is calculated as *B*_*0*_ – (percentage of bacteria-overlapped *C. roenbergensis* at the end of the second phase)×*B*_*0*_×2, assuming that each *C. roenbergensis* has ingested two bacteria. It is obvious that the fitting results for the third and fourth phases are consistent with less fitting errors than the results for the first and second phases. Mathematically this is because *k*_1_*B*_0_*t* and *k*_3_*t* are less than 1, which makes Taylor expansions converge quickly. The ingestion rate (*k*_*1*_) for each curve is 4-5 times slower in **Table 2** than its corresponding rate in **Table 1**.

**Table 2.**
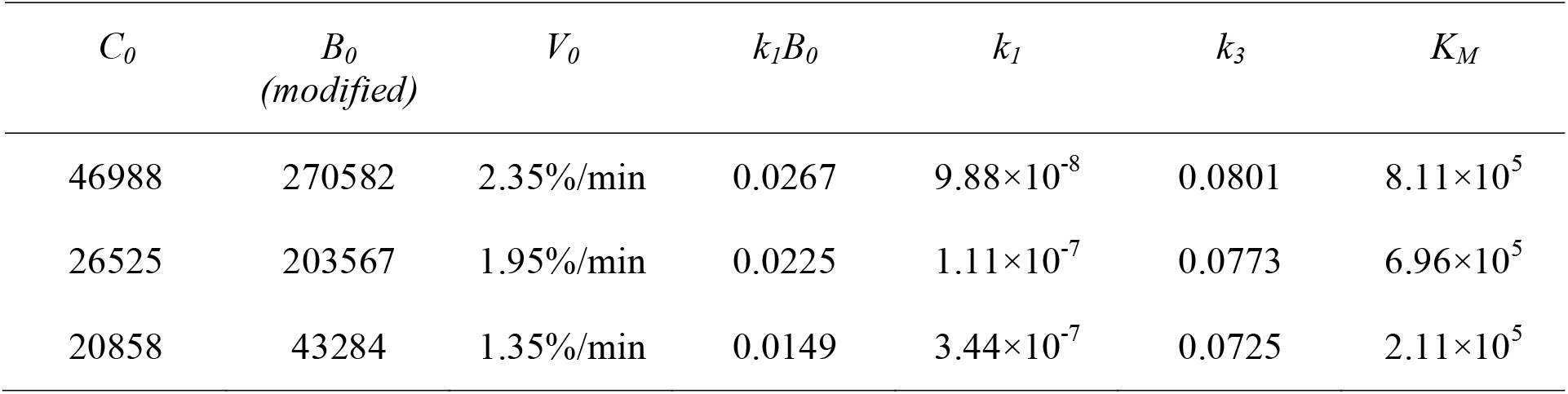
Fitting parameters for the third and fourth phases (16-35 minutes)

A common way to obtain reaction parameters such as *K*_*M*_ is to plot the initial reaction rate (*V*_*0*_) vs. substrate concentration (*S*) [13]. In this study, *V*_*0*_ is calculated as the initial slopes using the first two data points in **Figure 4 (a)** at *t*=0 and at 16 minutes respectively for the starved and fed cells then listed in **Tables 1** and **2** respectively, and the substrate concentrations are the total numbers of bacteria (*B*_*0*_) in each experiment in **Table 1** and **2**. Adapting the Lineweaver–Burk reciprocal plot will lead to a linear fit of the experimental data (**Figure 4 (b)**). The intercepting points of fitting curves with *x*-axis give the parameters − 1/*K*_*M*_, and the intercepting points with *y*-axis give the parameters 1/*V*_*max*_. *K*_*M*_ reflects the enzyme’s affinity for its substrates, whereas *V*_*max*_ represents the maximum rate at which the enzyme processes the substrate. **Table 3** lists the obtained parameters. The obtained *K*_*M*_ for starved cells in the initial period is on the same order of the estimation in **Table 1**, while the obtained *K*_*M*_ for fed cells in the later period is one order higher than the estimation in **Table 2**. Considering there are only three data points for regression and the unequal weighting of errors in Lineweaver–Burk reciprocal plot, this crude estimate is within reason range. In **Figure 4 (b)**, the two Lineweaver-Burk plots from the initial and later periods converged at the 1/*B*_*0*_ axis, which, in enzyme kinetics, is characteristic as a special case of noncompetitive inhibition [16]. In our specific case, *K*_*M*_ remains almost unchanged between the two periods, but *V*_*max*_ is reduced. These results suggest that the affinity between the predator and prey does not differ between the initial and later periods, i.e., the predator’s ability to capture the prey remains unchanged. However, certain inhibitory factors appear to slow down the predator’s efficiency in processing food during the later period, i.e. once the predators are fed, their digestion rate slows down. Previous observations have shown that starved cells graze at significantly higher rate than fed, exponential-growing cells [4]. However, no experiment has observed that starved cells slow down their grazing after feeding for a certain period of time, approximately 15 minutes. This might indicate that after the first rounds of feeding, the flagellate begins to shift their grazing dynamics to resemble those observed in fed, exponential growing cells. This transition has now been directly captured via two-photon fluorescence microscopic real-time videos.

**Table 3.** Fitting parameters from Lineweaver–Burk reciprocal plot.

| | $K_M$ | $V_{max}$ |
| --- | --- | --- |
| Initial Period | $6.02 \times 10^4$ | 24.0 |
| Later Period | $4.33 \times 10^4$ | 2.59 |

## Conclusions

In summary, this study presents a novel application of video-rate two-photon fluorescence microscopy to dynamically visualize and quantify the grazing behavior of the fast-swimming heterotrophic nanoflagellate *C. roenbergensis* on bacteria. By enabling real-time tracking of contact, ingestion, and digestion events in live, motile cells, this technique overcomes key limitations of traditional imaging methods and offers unprecedented insight into microbial predator-prey interactions. The observed transition from rapid to saturated feeding phases reflects physiological shifts from starvation to exponential growth. Compared to previous literature focus on individual flagellate, the quantitative modeling of grazing dynamics at the population level provides a foundation for linking microbial behavior to ecological function. This approach not only advances our understanding of microbial food webs but also establishes a broadly applicable platform for studying fast, transient interactions among diverse microbial species in aquatic environments. One area for future improvement is that the current video-rate microscopy lacks 3D tracking capability, i.e. it is difficult to change the focal plane of the microscope to observe fast moving objects in three-dimensional space. New optical microscopic technologies that can change the focal plane at high-speed, i.e. temporal focusing two-photon microscopy [17, 18], will provide new such capabilities and lead to more thorough investigations. Also, recent development in other optical microscopy techniques for 3D imaging, such as light sheet and light field microscopy [19, 20], could possibly overcome this obstacle.

## ACKNOWLEDGEMENTS

Research reported in this publication was supported by the National Science Foundation (NSF) under the award number 1429708 and 1205302, and the National Institute of General Medical Sciences (NIGMS) of National Institutes of Health (NIH) under the award number 1SC2GM103719 and R01GM129525. CX also received support from the Welch Foundation under Grant Number AH-2126-20220331. The content is solely the responsibility of the authors and does not necessarily represent the official views of the funding agencies.

## AUTHOR CONTRIBUTIONS

C.L. and C.X. conceived the concept and supervised the project. A.R., M.C., and Y.X. performed experiments and obtained and analyzed data with contributions from C.L. and C.X. C.L. and C.X. prepared the manuscript.

## SUPPORTING MATERIAL

Supporting materials can be found online.

