## Supplemental Methods for "Observing Grazing Behavior Transitions in *Cafeteria roenbergensis* with Video-Rate Two-Photon Microscopy"

In terms of kinetics the grazing behavior of *Cafeteria roenbergensis* is similar to enzyme-catalyzed biochemical reactions, in which *C. roenbergensis* (*C*) as an enzyme that takes the bacteria (*B*) as substrate and then degrade the bacteria into waste (*W*). The process will be

$$\boldsymbol{C + B \underset{k_1}{\overset{k_2}{\leftrightarrows}} C{\cdot}B \overset{k_3}{\rightarrow} C + W}$$

$k_1$ is the *C. roenbergensis* grazing rate, $k_2$ is the reverse reaction rate due to spitting out which was shown in some early papers [1], $k_3$ is the digestion rate. When this rection reaches an equilibrium state (steady-state assumption), the change of concentration of $C{\cdot}B$ with time $d\,[C{\cdot}B]\,/dt$ is zero, thus leading to

$$\frac{d[C{\cdot}B]}{dt} = k_1[C][B] - (k_2 + k_3)[C \cdot B] = 0 \qquad (1),$$

$$k_1[C][B] = (k_2 + k_3)[C \cdot B] \qquad (2).$$

The total *C. roenbergensis* concentration is a constant $C_0$ which is the concentration of free *C. roenbergensis* (that have not eaten) plus the bound *C. roenbergensis* concentration (that have engulfed bacteria but the bacteria is not fully digested yet.) $[C]_{tot} = C_0 = [C] + [\ C \cdot B]$. This can be rearranged as:

$$[C] = [C]_{tot} - [\ C \cdot B] = C_0 - [C \cdot B] \qquad (3).$$

Substituting $[C]$ in Eq. (2) from Eq. (3) results

$$k_1(C_0 - [\ C \cdot B])[B] = (k_2 + k_3)[C \cdot B]$$

$$k_1C_0[B] = (k_2 + k_3 + k_1[B])[C \cdot B]$$

$$\frac{[C \cdot B]}{C_0} = \frac{k_1[B]}{k_2 + k_3 + k_1[B]}$$

$$\frac{[C{\cdot}B]}{C_0} = \frac{[B]}{\frac{k_2+k_3}{k_1}+[B]} \qquad (4).$$

If we make a definition of

$$K_M = \frac{k_2+k_3}{k_1} \qquad (5),$$

the result is

$$\frac{[C{\cdot}B]}{C_0} = \frac{[B]}{K_M+[B]} \qquad (6),$$

and $K_M$ is the well-known Michaelis constant. The above derivations only consider steady state relations. The experimental data are time evolution of the concentration of $[C{\cdot}B]$. Nonlinear regression methods have been developed to extrapolate reaction parameters such as $K_M$ from such progressive measurements [2]. Since this is a simple reaction only involving one substrate, it is feasible to derive an analytical form for analysis. First, two coupled ordinary differential equations are laid out for describing the time evolution of both *C. roenbergensis* and bacteria concentrations:

$$\begin{cases} \frac{d[C\cdot B]}{dt} = k_1(C_0 - [C\cdot B])[B] - (k_2 + k_3)[C\cdot B] \\ \frac{d[B]}{dt} = -k_1 C_0[B] + k_2[C\cdot B] \end{cases} \quad (7).$$

These nonlinear equations are not analytically solvable. To simplify the spitting rate ($k_2$) of *C. roenbergensis* is zero assuming the *C. roenbergensis* does not spit bacteria considering they are the only food source. It has been discovered that *C. roenbergensis* stored all ingested particles, including both bacteria and plastic beads, for more than half hour [3], which indicates $k_2$ equals zero. Based upon this assumption Eq. (7) transform into

$$\begin{cases} \frac{d[C\cdot B]}{dt} = k_1(C_0 - [C\cdot B])[B] - k_3[C\cdot B] \\ \frac{d[B]}{dt} = -k_1 C_0[B] \end{cases} \quad (8).$$

Solving the second equation leads to

$$[B] = B_0 e^{-k_1 C_0 t} \quad (9),$$

where $B_0$ is the initial concentration of bacteria. Substituting [$C\cdot B$] in the first Eq. (8) from Eq. (9) results

$$\frac{d[C\cdot B]}{dt} = k_1 B_0 e^{-k_1 C_0 t}(C_0 - [C\cdot B]) - k_3[C\cdot B] \quad (10),$$

which can be rewritten as

$$\frac{d[C\cdot B]}{dt} = -(k_1 B_0 e^{-k_1 C_0 t} + k_3)[C\cdot B] + k_1 B_0 C_0 e^{-k_1 C_0 t} \quad (11).$$

The analytical solution of Eq. (11) is

$$[C\cdot B] = e^{-\int_0^t (k_1 B_0 e^{-k_1 C_0 t'} + k_3)dt'} \int_0^t k_1 B_0 C_0 e^{-k_1 C_0 t'} e^{\int_0^{t'} (k_1 B_0 e^{-k_1 C_0 t''} + k_3)dt''} dt' \quad (12).$$

Thus, Eq. (12) is the theoretical form for the progressive curve in **Figure 4**.

The first integration term gives

$$e^{\frac{B_0}{C_0}(e^{-k_1 C_0 t} - 1) - k_3 t} \quad (13).$$

And the second integration term gives

$$k_1 B_0 C_0 e^{\frac{B_0}{C_0}} \left[ \frac{1}{k_1 B_0} \left( e^{-\frac{B_0}{C_0} e^{-k_1 C_0 t} + k_3 t} - e^{-\frac{B_0}{C_0}} \right) - \frac{k_3}{k_1 B_0} \int_0^t e^{-\frac{B_0}{C_0} e^{-k_1 C_0 t'} + k_3 t'} dt' \right] \quad (14).$$

To further simplify Eq. (14), Taylor expansion is used and only the first two orders are considered to obtain

$$k_1 B_0 C_0 \left( t + \frac{(k_1 B_{0+} k_3)}{2}\ t^2 \right) \quad (15).$$

Eq. (13) can also be simplified with Taylor expansion as

$$e^{-(k_1 B_0 + k_3)t} \quad (16).$$

Together Eq. (12) becomes

$$[C \cdot B] = k_1 B_0 C_0 \left( t + \frac{(k_1 B_{0+} k_3)}{2}\ t^2 \right) e^{-(k_1 B_0 + k_3)t} \quad (17).$$

Since the progressive curve in **Figure 4** is the percentage of stained *C. roenbergensis* number over the total number of *C. roenbergensis*, this analytical form for this curve is

$$\frac{[C \cdot B]}{C_0} = k_1 B_0 \left( t + \frac{(k_1 B_{0+} k_3)}{2}\ t^2 \right) e^{-(k_1 B_0 + k_3)t} \quad (18).$$

Fitting data points with Eq. (18) will obtain the coefficients $k_1$ and $k_3$, thus, to estimate Michaelis constant as $K_M = \frac{k_3}{k_1}$.


1. Boenigk, J. and H. Arndt, *Particle Handling during Interception Feeding by Four Species of Heterotrophic Nanoflagellates.* Journal of Eukaryotic Microbiology, 2000. **47**(4): p. 350-358.
2. Duggleby, R.G., *[3] Analysis of enzyme progress curves by nonlinear regression*, in *Methods in Enzymology*. 1995, Academic Press. p. 61-90.
3. Boenigk, J., et al., *Confusing Selective Feeding with Differential Digestion in Bacterivorous Nanoflagellates.* Journal of Eukaryotic Microbiology, 2001. **48**(4): p. 425-432.
